# Protective effects of different doses of MitoQ separately and combined with trehalose on sperm function and antioxidative status of cryopreserved Markhoz goat semen

**DOI:** 10.1101/2022.08.22.504802

**Authors:** Ako Rezaei, Hamid Reza Bahmani, Shiva Mafakheri, Abbas Farshad, Parisa Nazari

**Affiliations:** Department of Animal Science, Kurdistan Agricultural and Natural Resources Research and Education Center, AREEO, Sanandaj, 6616936311, Iran; Department of Animal Science, Faculty of Agriculture, University of Kurdistan, Sanandaj 661715175, Iran

**Keywords:** MitoQ, semen cryopreservation, oxidative stress, mitochondrial activity, goat

## Abstract

The mitochondria-targeted antioxidant MitoQ has been regarded as an effective antioxidant agent against cryo-induced oxidative cellular damage. This study aimed to evaluate the use of different doses of MitoQ combined with trehalose to minimize mitochondrial impairment and oxidative stress during sperm cryopreservation of Markhoz goat. For this, semen collection was performed every 5 days from 5 bucks (10 ejaculates per buck). The ejaculates were pooled and then diluted in eight different Tris-based extenders as follows: no additives (control), 20, 200, 2000 nM of MitoQ (MT20, MT200, MT 2000, respectively), 150 mM of trehalose (Tr), MT20+Tr, MT200+Tr, MT2000+Tr. Each semen sample was frozen using a standard protocol, and sperm function and antioxidative status were evaluated after thawing. Results indicated higher total and progressive motility, acrosome and membrane integrity, superoxide dismutase, glutathione peroxidase, total antioxidant capacity, and lower DNA fragmentation and malondialdehyde in MT200+Tr than for all other groups except MT200; meanwhile, MT200 was also greater in these parameters than in the control group (P < 0.05). Furthermore, MT200 and MT200+Tr showed higher (P < 0.05) percentages of live cryopreserved sperm with high membrane mitochondrial potential than other groups. However, abnormality percentage and catalase activity of frozen-thawed sperm were not affected by treatments (P > 0.05). To conclude, we have found that supplementation of 200 nM MitoQ alone or in combination with 150 mM trehalose to semen extender improves the quality of cryopreserved sperm in goats, which is associated with enhanced antioxidant enzymatic defense and mitochondrial activity and reduced DNA fragmentation.

## 1. Introduction

Markhoz goat breed is faced a growing threat of extinction concurrent with diminished geographic distribution. The ex-situ programs aimed at cryopreserving germplasm are underway as a principal approach to setting national conservation priorities for Markhoz goats [11,30]. Sperm cryopreservation has been a standard procedure for the long-term preservation of male genetic materials, which can be transmitted by reproductive tools such as artificial insemination to offspring. The frozen-thawed sperm in the ruminants are exposed to structural and functional damage resulting from molecular element changes, including ATP-releasing nucleotides, lipids, proteins, etc. In the past decade, different cryoprotective agents, antioxidants and proteins have been incorporated into the freezing extender to prevent enhanced membrane fluidity-permeability, high reactive oxygen species (ROS) generation, acrosome integrity reduction, and the mitochondrial activity impairment of sperm after cryopreservation [22,43]. The ruminant sperm membrane has a higher unsaturated phospholipids/cholesterol ratio. Such composition is more sensitive to lipid peroxidation in the presence of ROS, which has less sperm cryo-resistance and, consequently, sperm cryo-survival. Besides sperm membrane cryodamage, oxidative stress diminishes membrane potential in mitochondria as the main organelle of endogenous ROS production [29]. The sperm mitochondrial membrane potential (MMP) is positively associated with motility and viability and negatively associated with DNA fragmentation [12]. Endogenous antioxidants are not adequately neutralized oxidation stress in the semen freezing process. Hence many studies have evaluated the cryoprotective effect of various exogenous antioxidants in semen extenders with target normalizing ROS levels through different oxidative pathways [5]. Exogenous antioxidants, including metabolites and ions less than 5 kDa size, are permitted to enter the external mitochondrial membrane, while none can penetrate the inner mitochondrial membrane [6,35]. Several mitochondria-targeted antioxidants have been designed to achieve direct antioxidant delivery in the inner mitochondrial membrane, which can provide higher concentrations at the ROS production site [33]. MitoQ is a recyclable mitochondrial-targeted antioxidant consisting of lipophilic triphenylphosphonium (TPP) cation as a carrier and ubiquinol moiety as an antioxidant. The TPP cation moiety (positive charges) on MitoQ can accumulate 100–1000 fold in the mitochondrial matrix (negative charges) driven by the membrane potential. The ubiquinol component of MitoQ in the mitochondrial electron transport chain is oxidized to ubiquinone and breaks chain free radical reaction, which is then restored by reduced at complex II [26,34]. Positive antioxidant effects of MitoQ on cardiac [10], hepatic [42], renal [17], and neuronal [39] mitochondrial status have been confirmed to protect against oxidative-associated diseases. Results from in vitro reports showed that the addition of MitoQ to a cooling medium protected mitochondrial function in rabbit heart [36], porcine [28], and rat [24] kidney after cold preservation. MitoQ could cross the rat blood-testis barrier and protect against rodent testicular injury by promoting the expression of mitochondrial dynamics proteins [44], activating antioxidant enzymes, and inhibiting apoptosis indicators [19]. The addition of MitoQ at various nanomolar concentrations in extender lowered the production of ROS and lipid peroxidation and improved sperm quality in frozen-thawed semen of catfish [13], human [21], buffalo [6], and rooster [25]. In buffalo semen, MitoQ indicated more cryo-tolerance than Resveratrol as a cytosolic antioxidant [38]. Improvement in sperm quality and antioxidant enzymatic activities was observed after the addition of 150 nM MitoQ to extender in cryopreserved ram semen; however, sperm mitochondrial function was not evaluated in this study [40].

On the other hand, trehalose, a non-permeable disaccharide, is used as a beneficial cryoprotectant agent in semen extenders to prevent cryo-shock damage by modifying osmotic pressure and interacting with sperm membrane structure [1,27]. The literature reports demonstrated the protection of cryopreservation-induced damage in the trehalose supplemented semen in small ruminant animals [22,29]. However, no studies have been reported on the cryo-protective effects of the combination of trehalose and mitochondria-targeted antioxidants.

The evidence above indicates that MitoQ is highly regarded for its efficient antioxidative performance on semen cryopreservation in most domestic ruminant species; while there is no statement about using MitoQ with goat sperm. In line with Markhoz breed conservation, we aimed to achieve the best doses of MitoQ separately and in combination with trehalose to propose an effective compound in the Tris-based extender of cryopreserved goat semen. Therefore, this study was carried out to evaluate the effects of different doses of MitoQ combined with trehalose on motility and velocity variables, quality characteristics (viability, abnormality, acrosome and membrane integrity, MMP and DNA fragmentation), and antioxidative status markers (superoxide dismutase [SOD], glutathione peroxidase [GPx], catalase, malondialdehyde [MDA], and total antioxidant capacity [TAC]) of frozen-thawed Markhoz goat semen.

## 2. Material and methods

### 2.1. Chemicals and ethics

Unless otherwise specified, all chemicals and reagents used were purchased from either Sigma (St. Louis, MO, USA) or Merck (Darmstadt, Germany). This study was approved by the Research Ethics Committee of the Animal Science Research Institute of Iran, with registration number ASRI901993 at 31.08.2020.

### 2.2. Animals, semen collection and preparation

Semen from 5 mature Markhoz bucks aged 2.5 to 3.5 years, with an average body weight of 68.2±3.6 kg, during the breeding season housed at Sanandaj Animal Husbandry Research Station [Kurdistan, Iran, (35.31 °N, 46.99 °E), day length: 11 h, humidity: 29 %, average day temperature: 16 °C] under uniform housing, feeding, and lighting conditions were used in this study. A total of 50 semen ejaculate samples (10 ejaculates for each buck) were collected at 5 days intervals from 9:00 to 10:00 am using the electroejaculation (Lane Manufacturing Inc., Denver, CO, USA) under sedation method according to the procedure described by Abril-Sánchez et al. [2]. Immediately after collection, ejaculates in each replication (n=10) with motility ≥75 %, normal morphology ≥ 85%, and 2×10^9^ sperm/mL were selected and pooled together to remove individual differences at 37°C.

### 2.2. Semen extender and frozen–thaw processing

The freezing extender was prepared with 3.786 gr Tris, 2.172 gr citric acid, 1 g fructose, 5 % glycerol (v/v), and 5 % (v/v) egg yolk in 100 ml double-distilled water at a pH of 6.8 and osmolality of 320 mOsm. The pre-evaluated pooled semen was split into eight equal aliquots and diluted in eight different Tris-based extenders at a concentration of 200×10^6^ sperm/mL as follows: no additives (control), 150 mM trehalose (Tr) (the best concentration of our unpublished data), 20 nM MitoQ (MedKoo Biosciences Inc., Morrisville, NC, USA) (MT20), 200 nM MitoQ (MT200), 2000 nM MitoQ (MT2000), 20 nM MitoQ + 150 mM trehalose (MT20+Tr), 200 nM MitoQ + 150 mM trehalose (MT200+Tr) and, 2000 nM MitoQ + 150 mM trehalose (MT2000+Tr). Extended samples were packaged in labeled 0.5 ml straws (IMV Technologies, l’Aigle, France), sealed with polyvinyl chloride powder, and then gradually chilled at 5 °C for 5 h. The straws were placed horizontally 4 cm above the liquid nitrogen surface for 15 min and then immersed directly in a cryobiological (−196 °C) container until used. Stored straws were thawed individually at 37 °C for 30 s in a water bath and immediately evaluated for sperm characteristics as follows;

### 2.3. Evaluation of sperm motion characteristics

For sperm motility and velocity evaluation, a 5 μl semen sample per replicate were placed in a pre-warmed chamber slide (Leja 4, Leja Products, Luzernestraat B.V., Holland) and immediately covered with glass coverslips. At least 10 microscopic fields to include 300 sperms for each sample were analyzed using a computer-assisted sperm analysis (CASA) system (IVOS version 12; Hamilton-Thorne Biosciences, MA, USA) equipped with a digital video camera (Samsung, SDC-313B, Korea) attached to phase-contrast microscope (Nikon, Eclipse E200, Japan). The CASA-derived motility characteristics studied were total motility (%); progressive motility (%); VAP: average path velocity (μm/s); VSL: straight linear velocity (μm/s); VCL: curvilinear velocity (μm/s); STR [VSL/VAP]: straightness (%); LIN [VSL/VCL]: linearity (%); ALH: amplitude of lateral head displacement (μm); BCF: beat cross frequency (Hz).

### 2.4. Evaluation of sperm quality characteristics

#### 2.3.1. Sperm viability

The eosin-nigrosin stain procedure was used to evaluate sperm viability [14]. In brief, 10 ml of sperm sample was mixed with stain (eosin Y 0.67 g, nigrosin 10 g, NaCl 0.9 g dissolved in 100 ml distilled water) and smeared on a pre-warmed slide. The smears were air-dried and observed 200 sperm under the bright field microscope at 400× magnification. Sperm viability was obtained by the percentage of non-stained head sperm/total sperms.

#### 2.3.2. Sperm abnormality

The Hancock solution (150 ml PBS buffer, 62.5 ml of 37% formalin, 150 ml sodium saline solution, and 500 ml double-distilled water) was used to evaluate sperm morphological abnormalities [31]. This was performed by mixing 50 μl of each semen sample with 1 ml of Hancock solution in an Eppendorf tube. Then, 10 μl of mixture sample was loaded on a pre-warmed slide and covered with a coverslip. The percentage of sperm cells with abnormal heads and/or tails was recorded by counting 200 sperms using a phase-contrast microscope (1000× magnification, oil immersion).

#### 2.3.3. Sperm membrane integrity

The hypo-osmotic swelling test (HOST) was used for sperm membranes evaluation [16]. With this procedure, 5 μl of semen sample was mixed with 500 μl of 1% formal citrate (9 g fructose + 4.9 g sodium citrate per liter of distilled water) and incubated at 37 °C for 60 min. Then, 100 μl of the mixture was loaded on a pre-warmed slide covered with a coverslip. A total of 200 sperm from at least 10 microscopic fields was counted for the percentage of sperms with swollen and curled tails under a phase-contrast microscope (400 × magnification).

#### 2.3.4. Sperm acrosome integrity

The evaluation of sperm acrosome integrity was carried out using a modification of the buffered formalin-citrate solution described by Ahmad et al. [4].In brief, 100 μl of semen sample was fixed with 500 μL of formal citrate (96 ml 2.9 tri-sodium citrate dehydrate with 4 ml of 37% formaldehyde) on a slide covered with a coverslip. After drying in air, 200 sperms in at least 10 microscopic fields were counted for the percentage of sperms acrosome with normal apical ridge under a phase-contrast microscope (1000× magnification).

#### 2.3.4. Sperm MMP

The sperm MMP was evaluated using two fluorescent stains containing Rhodamine-123 (R123; Invitrogen TM, Eugene, OR, USA) and propidium iodide (PI) by flow-cytometric analysis. In brief, 300 ml of diluted semen sample was incubated with 10 μl Rhodamine (0.01 mg/ml) for 20 minutes at room temperature (RT) in the dark. The processed semen samples were centrifuged for 3 min at 500 g and resuspended in 500 μl Tris buffer. Next, 10 μl of PI (1 mg/ml) was added to the mixture and analyzed with a FACS Calibur Flow cytometer (Becton Dickinson, San Jose, USA). The percentage of sperm with active functional mitochondria was characterised by positive for R123 and negative for PI. At least 10000 events were recorded per sample [7].

#### 2.3.5. Sperm chromatin dispersion

The sperm chromatin dispersion (SCD) test was performed to evaluate sperm DNA fragmentation according to the procedure of Fernandez et al. [15]. In brief, 150 μl of 65% agarose were loaded on a slide, covered with a coverslip, and stored at 4 °C for 5 min. After removing the coverslip, 30 μl of semen sample was mixed with 70 μl of low melting point agarose (0.7% in PBS) and transferred on the agarose-covered slide, covered with a coverslip quickly, and allowed to air dry. The coverslip was removed, and the slide was immersed in 0.08 N HCl solution for 7 min at 37 C in the dark. The slide was gently placed in lysing solution (0.4 M Tris, 0.8 M DDT, 1% SDS, 2 M NaCl, and 50 mM EDTA, pH 7.5) for 25 min at RT. The slide was washed with distilled water for 5 min, dehydrated in 70%, 90% and 100% ethanol, respectively, for 2 min. After drying in air, the halo surface/whole nucleoid surface (HS/WNS) ratio in sperm stained with 0.2 mg/mL ethidium bromide was investigated using an inverted fluorescence microscope (Olympus CKX53, Tokyo, Japan) as presented in Fig. 1.

**Fig. 1.**
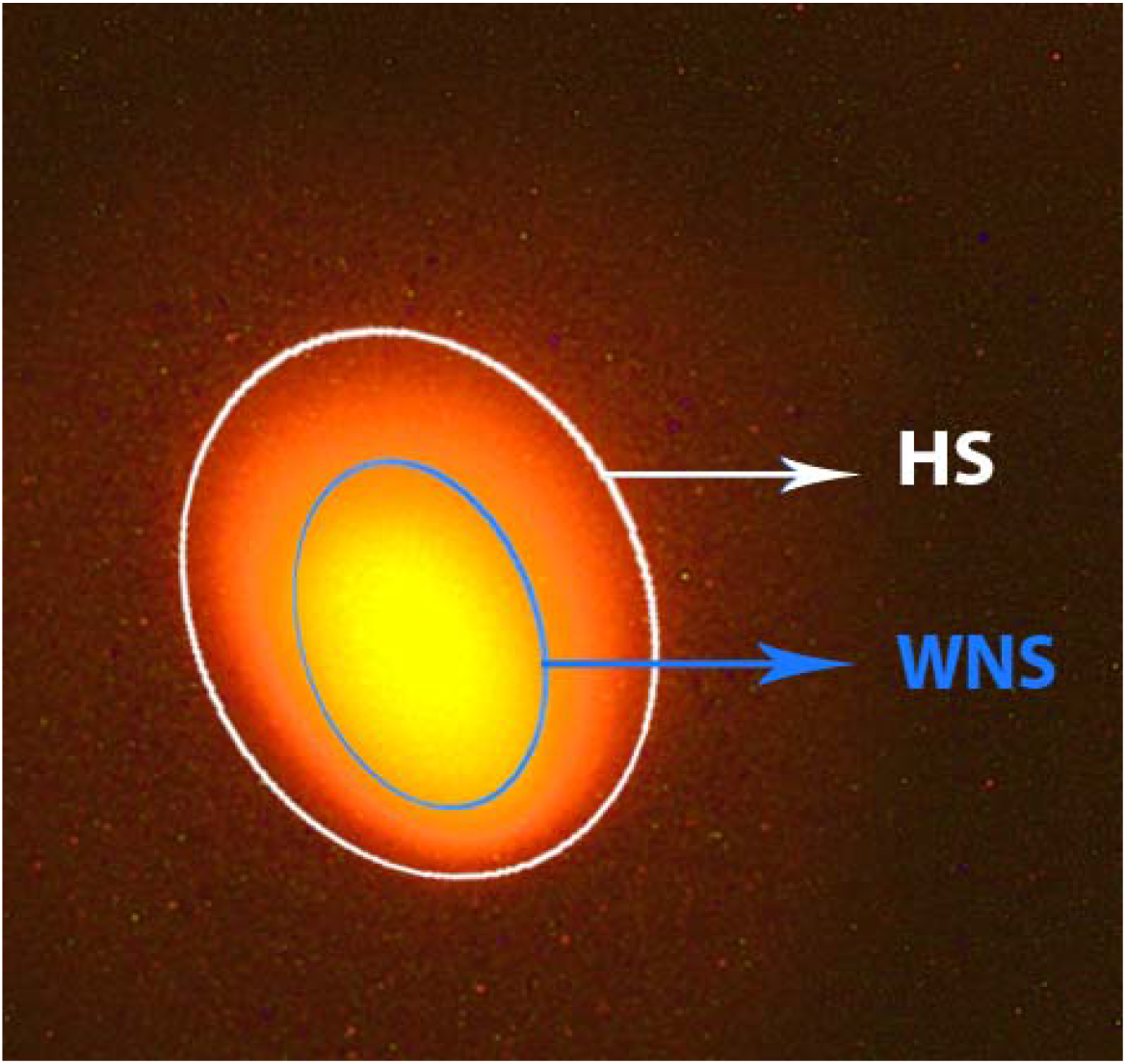
Fluorescence image of sperm stained by ethidium bromide. Relative halo size was calculated by the division of halo surface (HS) to whole nucleoid surface (WNS).

### 2.4. Evaluation of sperm antioxidative characteristics

For antioxidant indices assays, the 500 μl of thawed semen sample was centrifuged (500 g for 20 min), and the sperm was collected, washed with PBS, and recentrifuged to obtain a concentration of 2×10^8^ sperm/ml [8].

#### 2.4.1. Sperm MDA concentration

Evaluation of MDA concentration in the sperm sample as a measure for lipid peroxidation was determined by Thiobarbituric acid (TBA) reaction as described by Greg et al. [18]. The first, 1 ml of prepared sperm was added to 2 ml of TBA-trichloroacetic acid (TCA) solution (0.375% w/v TBA, 15% w/v TCA, and 0.25 N HCl). The mixture was boiled in a water bath at 100 °C for 15 min and then cooled at RT. Then after centrifugation at 1200 g for 15 min, the absorbance of the supernatant was measured by a Hitachi U-2001 spectrophotometer (Tokyo, Japan) at 535 nm. Finally, the MDA concentration was calculated using the specific absorbance coefficient (1.56×10^5^/mol/cm^3^) and expressed as nmol/ml.

#### 2.4.2. Sperm antioxidant enzymes (SOD, GPx and catalase)

The activities of SOD and GPx in the sperm samples were determined using Ransel and Ransod kits, respectively, from Randox Laboratories (Crumlin, UK) following the manufacturers′ instructions. The SOD assay is based on the inhibition of red formazan dye generated by reaction of 2-(4-iodophenyl)-3-(4-nitrophenol)-5-phenyltetrazolium chloride with the superoxide radicals generated by xanthine and xanthine oxidase. SOD activity was determined by monitoring the 50% inhibition of the formazan dye formation at 505 nm and expressed as U/mg protein. The GPx activity was measured following the oxidation of the reduced form NADPH with glutathione reductase and reduced glutathione using a cumene hydroperoxide probe. Next, GPX activity was quantified by absorbance of NADPH oxidation at 340 nm and expressed as U/mg protein. Catalase activity in the sperm sample was determined through the decomposition rate of hydrogen peroxide into O2 and H_2_O_2_ according to the procedure of Aebi [3]. For this assay, 20 μl of prepared sperm was mixed to reaction solution (830 μl of 50 mM potassium phosphate buffer and 150 μl of 30 mM H_2_O_2_, pH 7.2). The mixture absorbance was measured for 3 min at 240 nm, and catalase activity was expressed as U/mg protein. Spectrophotometric analysis of enzymatic activities was performed by an Alcyon 300 autoanalyzer (Abbott, IL, US). Total protein was determined by the Bradford Protein Assay Kit (Bio-Rad, Hercules, CA, USA) with bovine serum albumin as a standard and normalized all enzymatic data.

#### 2.4.3. Sperm TAC concentration

The TAC assay was performed in the semen sample using a colorimetric method by a commercial kit (Randox, Crumlin, UK) per the manufacturer’s instructions. The principle of the assay was based on the ability of sample antioxidants to inhibit oxidation of the ABTS (2,2′-azinodisulfinate 3-ethylbenztiazoline) to radical cation (ABTS^+^), which was compared with a Trolox (reference standard antioxidant). The percentage inhibition of absorbance of blue-green ABTS+ in prepared samples was measured spectrophotometrically (autoanalyzer; Abbott, Alcyon 300, IL, US) at 600 nm.

### 2.5. Statistical analysis

Data normality and homogeneity of variances were verified using the Shapiro–Wilk and Levene’s tests, respectively. Data were statically analyzed under a completely randomized design using the General Linear Model procedure of SAS version 9.1 (SAS Institute, Cary, NC, USA). An analysis of orthogonal contrasts was used to evaluate differences among groups. Group means were compared using Tukey’s multiple range tests at P < 0.05 and stated as mean ± standard error of the mean.

## 3. Results

### 3.1. Sperm motion characteristics

The CASA-derived motility and velocity characteristics are shown in Table 1. The percentage of total and progressive motility were significantly higher (P < 0.05) for MT200+Tr than for all other groups except MT200. The MT200 group was significantly greater (P < 0.05) in total and progressive motility percentage compared to MT20, MT2000, MT2000+Tr, Tr, and control groups. Also, higher (P < 0.05) total and progressive motility percentages were found in MT20 and MT20+Tr than in control. The VAP of sperm was greater (P < 0.05) in the MT200 and MT200+Tr semen samples compared to the MT2000 and control semen samples. MT200+Tr group had greater VSL of sperm than all other groups except for MT200 and MT20+Tr. The MT200 and MT20+Tr groups also showed higher VSL of sperm than MT20, MT2000, MT2000+Tr, Tr, and control groups (P < 0.05). However, no differences occurred in the VCL, STR, ALH, LIN, and BCF of sperm samples between the groups (p > 0.05).

**Table 1.** Mean (±SEM) sperm motion characteristics in frozen–thawed Markhoz goat semen with different doses of MitoQ (20, 200 and 2000 nM) combined with trehalose (150 mM).

| Parameter | Control | MitoQ (nM) |  |  | Trehalose (150 mM) | MitoQ (nM) + Trehalose (150 mM) |  |  |
| --- | --- | --- | --- | --- | --- | --- | --- | --- |
|  |  | MT<sub>20</sub> | MT<sub>200</sub> | MT<sub>2000</sub> | Tr | MT<sub>20</sub>+Tr | MT<sub>200</sub>+Tr | MT<sub>2000</sub>+Tr |
| TM (%) | 43.62±1.22<sup>d</sup> | 48.14±1.37<sup>c</sup> | 52.00±1.01<sup>ab</sup> | 43.25±1.26<sup>d</sup> | 46.25±1.23<sup>cd</sup> | 48.57±0.89<sup>bc</sup> | 53.75±1.48<sup>a</sup> | 45.00±0.83<sup>cd</sup> |
| PM (%) | 39.37±0.70<sup>d</sup> | 42.00±0.81<sup>c</sup> | 44.87±0.66<sup>ab</sup> | 39.00±0.73<sup>d</sup> | 41.25±1.11<sup>cd</sup> | 42.75±1.10<sup>bc</sup> | 46.12±0.93<sup>a</sup> | 40.37±0.86<sup>cd</sup> |
| VAP(μm/s) | 60.18±1.57<sup>b</sup> | 63.13±1.71<sup>ab</sup> | 66.06±1.04<sup>a</sup> | 60.03±1.44<sup>b</sup> | 63.76±1.58<sup>ab</sup> | 64.42±1.80<sup>ab</sup> | 66.53±1.26<sup>a</sup> | 61.09±1.45<sup>ab</sup> |
| VSL (μm/s) | 36.51±0.75<sup>d</sup> | 38.28±0.79<sup>bcd</sup> | 40.49±0.64<sup>ab</sup> | 36.08±0.87<sup>d</sup> | 39.11±0.77<sup>bc</sup> | 39.99±0.96<sup>ab</sup> | 41.74±0.65<sup>a</sup> | 37.20±0.72<sup>cd</sup> |
| VCL(μm/s) | 103.84±1.85 | 106.89±2.36 | 109.89±2.52 | 103.41±2.53 | 106.82±2.65 | 107.72±2.64 | 110.76±2.37 | 105.65±2.39 |
| STR (%) | 60.84±1.36 | 60.99±2.15 | 61.38±1.24 | 60.52±2.12 | 61.72±2.41 | 62.57±2.34 | 62.90±1.51 | 61.03±1.29 |
| ALH (μm) | 4.70±0.25 | 5.11±0.36 | 5.22±0.23 | 4.69±0.18 | 5.00±0.13 | 5.17±0.23 | 5.36±0.24 | 4.79±0.27 |
| LIN (%) | 35.23±0.92 | 35.89±0.82 | 36.98±0.99 | 35.10±1.13 | 36.73±1.09 | 37.21±0.95 | 37.72±0.62 | 35.31±0.91 |
| BCF (Hz) | 11.05±0.60 | 11.66±0.73 | 12.28±0.31 | 10.83±0.38 | 11.48±0.49 | 12.02±0.55 | 12.26±0.49 | 11.15±0.46 |

<sup>a-b-c-d</sup> Values within rows with different letters are significantly different ( $P < 0.05$ ). Tr: 150 mM trehalose; MT20: 20 nM MitoQ; MT200: 200 nM MitoQ; MT2000: 2000 nM MitoQ. TM: total motility; PM: progressive motility; VAP: average path velocity; VSL: straight linear velocity; VCL: curvilinear velocity; STR: sperm track straightness; LIN: linearity; ALH: amplitude of lateral head displacement; BCF: beat cross frequency.

### 3.2. Sperm quality characteristics

Table 2 presents the data related to sperm quality characteristics. MT200 and MT200+Tr revealed higher (P < 0.05) percentages of live cryopreserved sperm than other groups. Moreover, higher (P < 0.05) percentages of acrosome and membrane integrity were recorded for MT200+Tr than for all other groups except MT200. The MT200 group showed higher acrosome integrity percentages than MT2000 and control groups (P < 0.05), and higher membrane integrity percentages than MT20, MT2000, MT2000+Tr, and control groups. However, no significant differences appeared between extenders for abnormal sperm percentage (P > 0.05).

**Table 2.** Mean (±SEM) sperm quality characteristics in frozen–thawed Markhoz goat semen with different doses of MitoQ (20, 200 and 2000

| Parameter | Control | MitoQ (nM) |  |  | Trehalose (150mM) | MitoQ (nM) + Trehalose (150 mM) |  |  |
| --- | --- | --- | --- | --- | --- | --- | --- | --- |
|  |  | MT<sub>20</sub> | MT<sub>200</sub> | MT<sub>2000</sub> | Tr | MT<sub>20</sub>+Tr | MT<sub>200</sub>+Tr | MT<sub>2000</sub>+Tr |
| VS (%) | 50.71±1.24<sup>c</sup> | 53.14±0.87<sup>bc</sup> | 59.12±1.00<sup>a</sup> | 49.75±0.88<sup>c</sup> | 52.87±1.37<sup>bc</sup> | 54.71±0.80<sup>b</sup> | 60.50±1.21<sup>a</sup> | 51.37±1.36<sup>bc</sup> |
| TA (%) | 25.54±1.70 | 23.96±1.72 | 23.27±0.97 | 26.07±1.13 | 24.28±1.26 | 23.53±0.88 | 22.58±0.68 | 24.88±0.96 |
| MI (%) | 42.12±0.85<sup>d</sup> | 43.86±1.13<sup>cd</sup> | 49.14±1.22<sup>ab</sup> | 41.62±1.03<sup>d</sup> | 44.00±0.77<sup>cd</sup> | 46.87±0.98<sup>bc</sup> | 52.37±1.43<sup>a</sup> | 43.75±0.97<sup>d</sup> |
| AI (%) | 57.14±0.77<sup>c</sup> | 58.00±0.69<sup>bc</sup> | 59.71±0.81<sup>ab</sup> | 56.87±0.51<sup>c</sup> | 57.75±0.93<sup>bc</sup> | 58.12±0.83<sup>bc</sup> | 60.75±1.18<sup>a</sup> | 57.62±0.70<sup>bc</sup> |

nM) combined with trehalose (150 mM)
<sup>a-b-c-d</sup> Values within rows with different letters are significantly different ( $P < 0.05$ ). Tr: 150 mM trehalose; MT20: 20 nM MitoQ; MT200: 200 nM MitoQ; MT2000: 2000 nM MitoQ. VS: viable sperm; TM: total abnormality; AI: acrosome integrity; MI: membrane integrity.

In Figs. 2 and 3, the observation of SCD test indicates that HS/WNS ratio in MT200+Tr was higher (P < 0.05) than in all other groups except MT200. The MT200 group also showed an elevation (P < 0.05) of the HS/WNS ratio compared with MT2000, MT2000+Tr, Tr, and control groups. As depicted in Fig. 4, the percentage of sperm MMP was higher (P < 0.05) in MT200 and MT200+Tr compared to other groups.

**Fig. 2.**
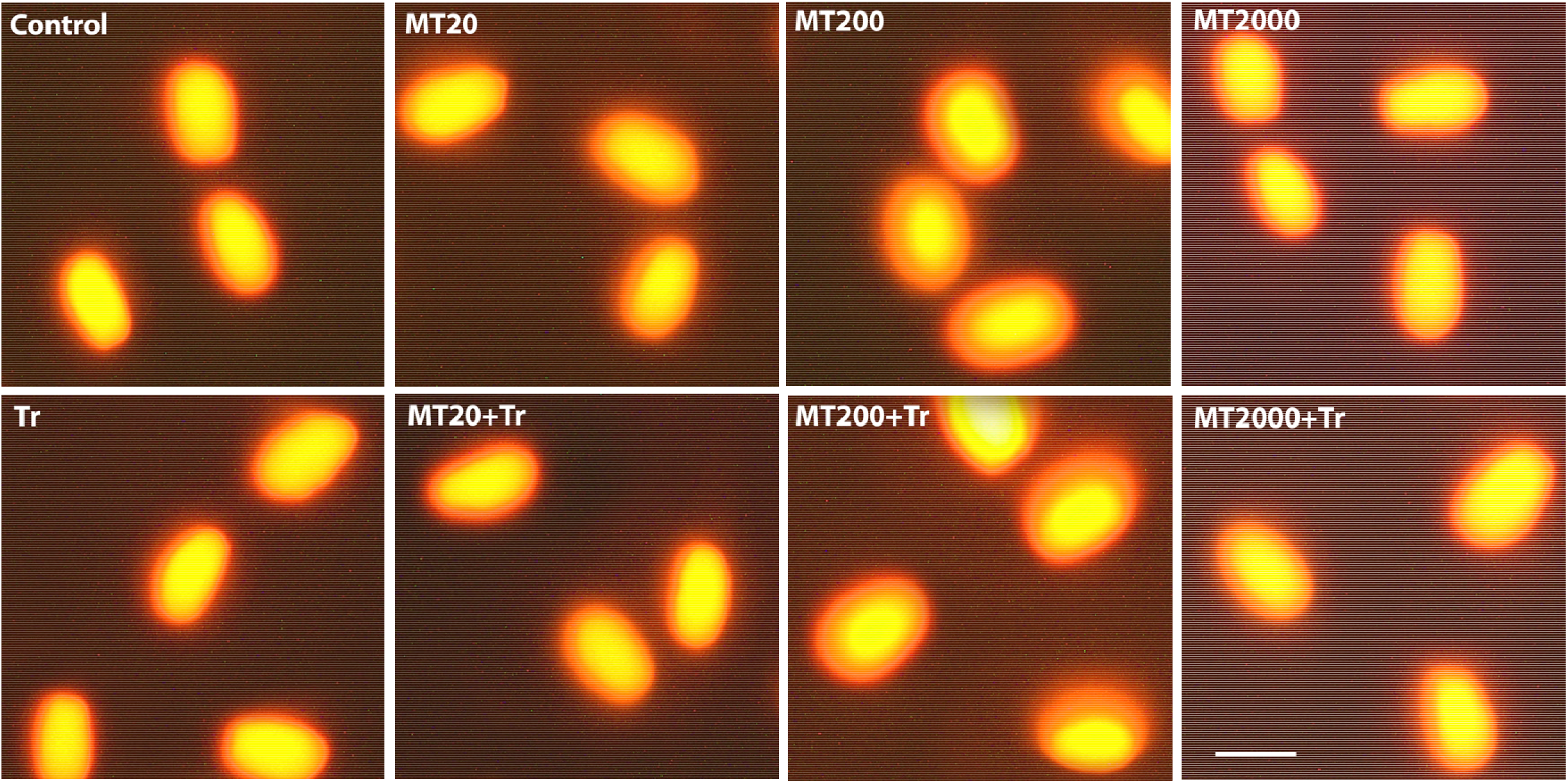
Representative images of sperm DNA fragmentation in frozen–thawed Markhoz goat semen with different doses of MitoQ (20, 200 and 2000 nM) combined with trehalose (150 mM). Tr: 150 mM trehalose; MT20: 20 nM MitoQ; MT200: 200 nM MitoQ; MT2000: 2000 nM MitoQ.

**Fig. 3.**
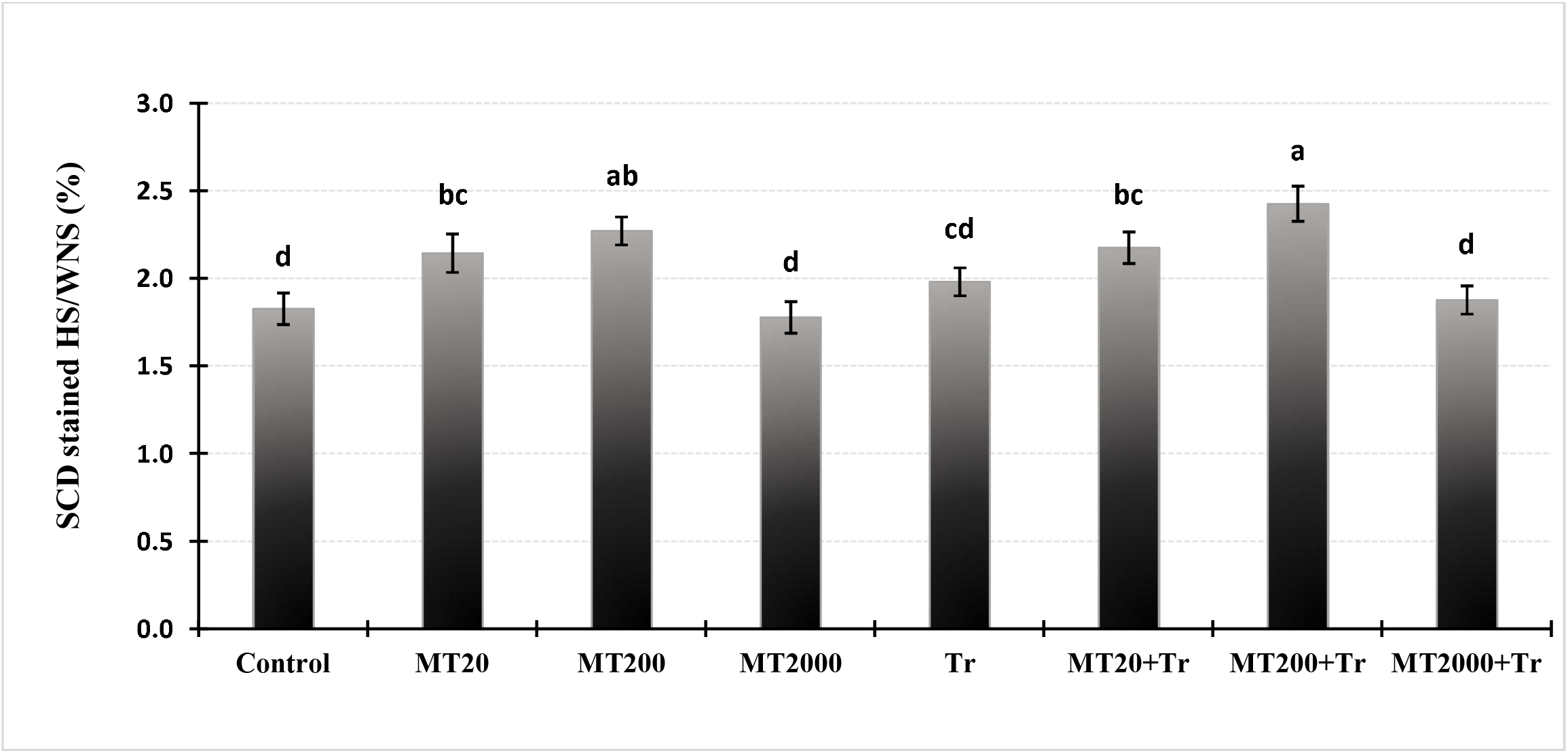
Sperm chromatin dispersion in frozen–thawed Markhoz goat semen with different doses of MitoQ (20, 200 and 2000 nM) combined with trehalose (150 mM). Error bars indicate standard errors of the mean. ^a-b-c-d^ columns with different letters are significantly different (P < 0.05). Tr: 150 mM trehalose; MT20: 20 nM MitoQ; MT200: 200 nM MitoQ; MT2000: 2000 nM MitoQ; SCD: sperm chromatin dispersion; HS/WNS: halo surface/whole nucleoid surface.

**Fig. 4.**
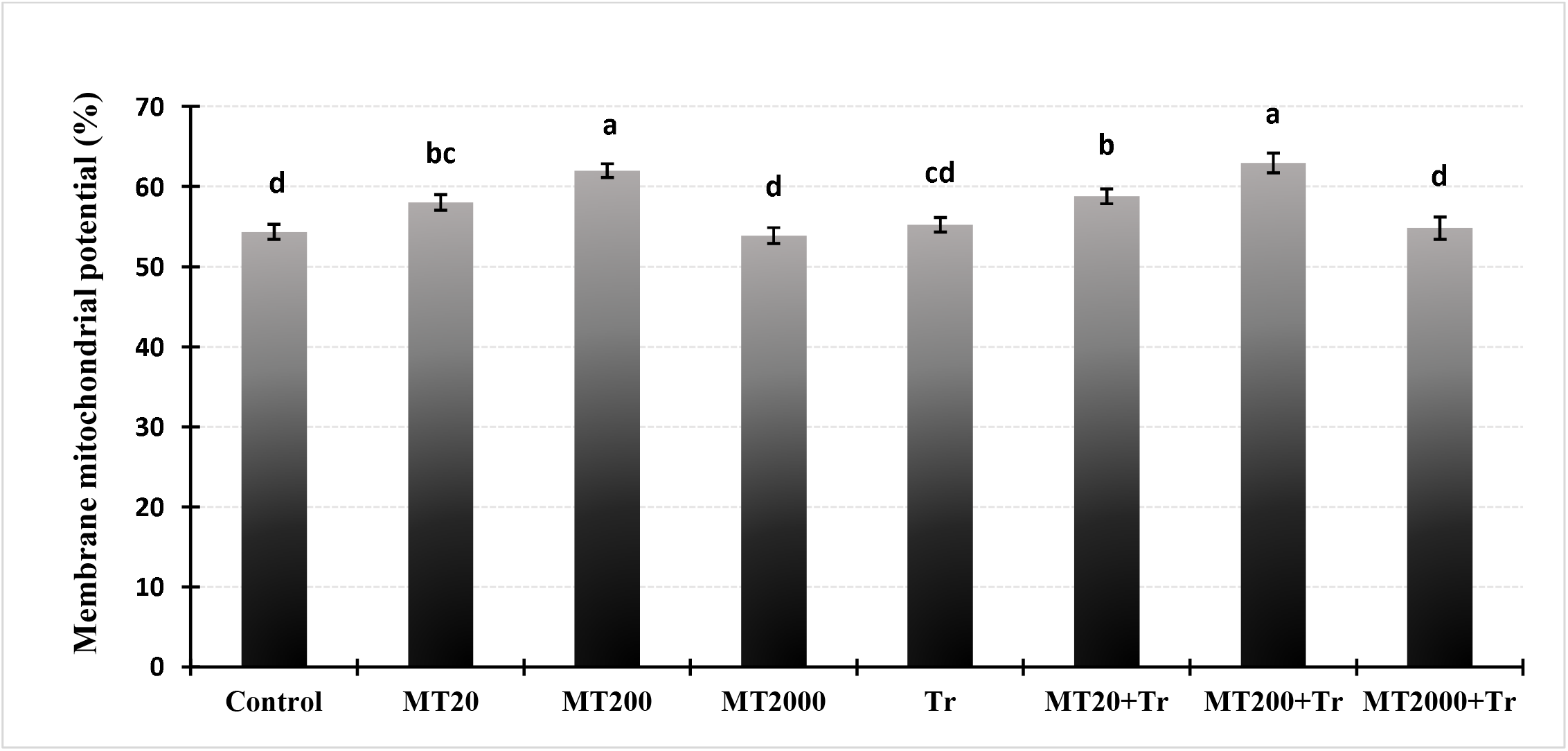
Sperm membrane mitochondrial potential in frozen–thawed Markhoz goat semen with different doses of MitoQ (20, 200 and 2000 nM) combined with trehalose (150 mM). Error bars indicate standard errors of the mean. ^a-b-c-d^ columns with different letters are significantly different (P < 0.05). Tr: 150 mM trehalose; MT20: 20 nM MitoQ; MT200: 200 nM MitoQ; MT2000: 2000 nM MitoQ.

### 3.2. Sperm antioxidative characteristics

Table 3 presents the mean concentration of MDA, SOD, GPx, CAT, and TAC in cryopreserved sperm. The sperm MDA concentration in the MT200+Tr and MT200 sperm samples was lesser (P < 0.05) compared to the MT2000, MT2000+Tr, Tr, and control sperm samples. Furthermore, the SOD activity of cryopreserved sperm was higher (P < 0.05) in MT200+Tr compared with all other groups except for MT200. The MT200 group also indicated higher sperm SOD activity than MT2000, MT2000+Tr, Tr, and control groups (P < 0.05). Results of the sperm GPx activity indicated higher (P < 0.05) for MT200+Tr than for MT2000, MT2000+Tr, Tr, and control groups. In addition, regarding sperm TAC concentration, MT200+Tr showed (P < 0.05) higher than all other groups except for MT200. The MT200 group also showed higher sperm TAC concentration than MT2000, MT2000+Tr, Tr, and control groups (P < 0.05). Moreover, no differences (P > 0.05) occurred in the catalase activity of sperm cryopreserved between the groups.

**Table 3.**
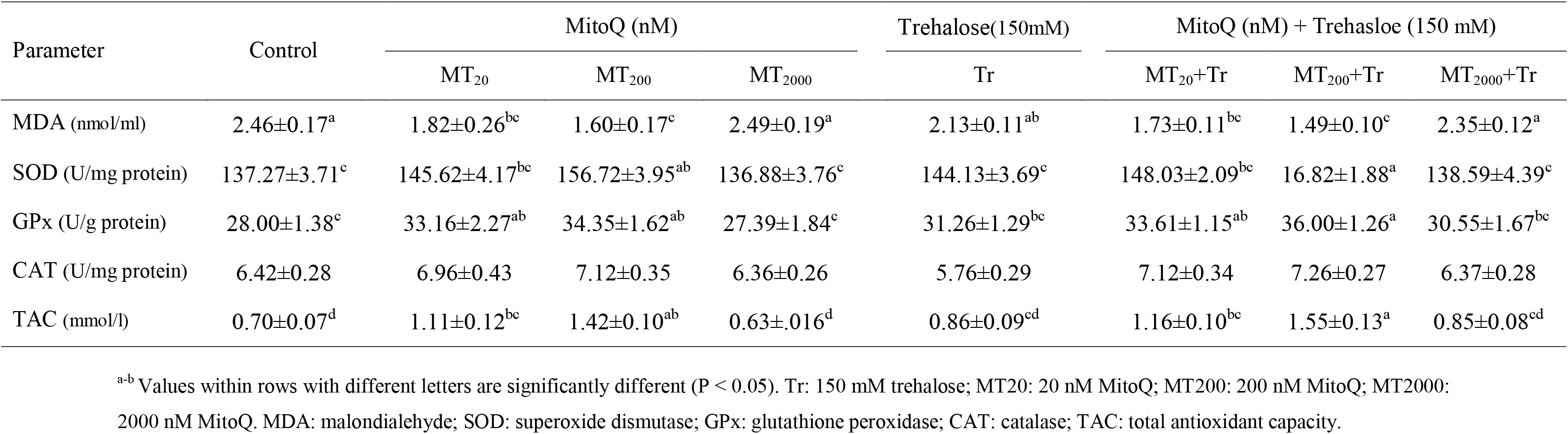
Mean (±SEM) sperm antioxidative characteristics in frozen–thawed Markhoz goat semen with different doses of MitoQ (20, 200 and 2000 nM) combined with trehalose (150 mM).

## 4. Discussion

To the best of our knowledge, this study is the first to report the MitoQ effects on semen cryopreservation in goats, which could be helpful for bio-conservation of the Markhoz breed. The expected adverse consequences of cryopreservation on sperm quality are firstly associated with plasma membrane disruption and then mitochondrial dysfunction, as reported by Peris-Frau et al. [29]. Based on this, we use trehalose as an osmoprotectant of membrane structure and MitoQ as an effective antioxidant targeted to mitochondria in the Tris-based extender with the aim of improving cryopreserved goat sperm function characteristics. In the present study, total motility, progressive motility, VAP, and VSL of post-thawed sperm were significantly better in MT200 and MT200+Tr groups, and other CASA-derived motility characteristics were the highest with MT200+Tr in the extender. This is congruent with previous studies, which have demonstrated the extender containing the concentration of 200 nM in human [21] and buffalo [38], and 150 nM in ram [40] have better sperm motility after cryopreservation. MitoQ supports ATP synthesis to maintain sperm motility through the electrons exchange of the redox cycle in the mitochondrial electron transport chain [37]. Furthermore, sperm proteins are susceptible to high ROS levels, which could negatively affect sperm flagellar movement. Similarly, a recent study showed that addition MitoQ to the human vitrification medium leaves activation of adenylate kinase 2 in sperm flagella that were necessary to sperm normal motility [20].

In our study, the extenders with 200 nM of MitoQ presented better post-thawed sperm quality characteristics such as viability, acrosome and membrane integrity, MMP and DNA fragmentation, which is in agreement with previous studies [6,13,25,38,41]. This preserving sperm quality can be related to the ability of MitoQ to directly lowed ROS levels in mitochondria. Consistently with this, MT200 groups in the present study showed that the variation trends of sperm quality coincided with improving sperm antioxidant indexes (SOD, GPx, TAC, and MDA levels). ROS plays vital roles in chromatin remodeling, hyperactivation, acrosome reaction, and oocyte fusion of sperm. Following sperm cryopreservation, the superoxide anion as the principal mitochondrial ROS increases, which promotes lipid peroxidation and ultimately impairs sperm function [37]. MitoQ has been able to diminish superoxide anions in frozen-thawed sperm, which could be explained by the enhanced mitochondrial antioxidant enzymes, especially SOD [6]. SOD and GPx enzymes act as the primary intracellular defenses against stress oxidative by detoxifying ROS. SOD dismutase superoxide anion to hydrogen peroxide and oxygen, and GPx also catalyzes hydrogen peroxide as SOD product, which can reduce lipid peroxidation and oxidative stress. In this context, an earlier study has demonstrated that the MitoQ addition to the ram sperm freezing extenders increases SOD and GPX and reduces ROS and MDA levels [40]. Therefore, it seems likely that MitoQ used in our study reduced mitochondrial ROS by replenishing antioxidant capacity and increasing SOD and GPx, which resulted in lower MDA as a lipid peroxidation product in frozen-thawed sperm goats.

An intact sperm plasma membrane is a prerequisite for acrosome reaction initiation and oocyte fusion. In the present study, the higher plasma membrane integrity in MT200+Tr could be explained by MitoQ effects on ROS reduction to prevent membrane lipid peroxidation [37]. Another possibility is that trehalose modulated membrane fluidity by interaction with the bilayer surface of the membrane [1]. Acrosome has a specialized structure consisting of membranes and proteins sensitive to high ROS levels. Apart from lipid peroxidation, oxidation of acrosomal proteins might also contribute to acrosomal damage after cryopreservation. During human sperm vitrification, MitoQ prevents alteration of proteins such as actin-like protein 7B and talin-1 localized in the acrosomal region, leading to maintain acrosome integrity [20], which is in accord with our result study.

Besides its bioenergetic function and ROS generation, sperm mitochondria are also involved in calcium homoeostasis and the intrinsic apoptosis pathway. Sperm activity is dependent on the integrity and functionality of mitochondria. MMP, as an electrostatic potential, is produced by creating an electrochemical gradient across the intermembrane space of mitochondria. MMP has been considered a predictor of sperm quality, which reflects the energy status, permeability transition, and respiratory chain activity in mitochondria [12]. During cryopreservation, MMP reduced by increasing permeability leads to the release of pro-apoptotic factors from the mitochondria, resulting in less sperm viability [23]. We found that the sperms treated with 200 nM MitoQ had higher MMP and viability than other groups. The obtained results agree with the previous study in buffalo cryopreserved semen [38], which indicated 200 nM MitoQ increased live sperms and MMP, and decreased apoptotic and necrotic sperms. Similar studies demonstrated that the MitoQ supplemented with semen freezing extender had greater live sperms with active mitochondria in catfish [13], human [21], and buffalo [6]. On the contrary, a recent report by Câmara et al. [9] showed that 0.2 to 20 nM MitoQ did not affect sperm quality and mitochondrial performance after bull semen cryopreservation, which might be due to the use of low concentrations of MitoQ in this study.

Our results have shown that extender groups containing 200 nM MitoQ lower dispersion of sperm chromatin as an indicator of DNA fragmentation accompanied by increased MMP and antioxidative status after semen cryopreservation. These findings align with those of previous studies, which revealed that MitoQ could alleviate biomarker of oxidative damage to DNA in mouse testis [44] and rat brain [39]. This DNA damage protection might be attributed to the effects of ubiquinol on DNA methylation patterns, as suggested by Schmelzer and Döring [32]. Indeed, it has been established that DNA damage is associated with ROS overproduced by mitochondria [12]. Therefore, in the present study we can suggest that the lower DNA fragmentation in MitoQ treatment may be due to decreased leakage of ROS by improving MMP and antioxidant defense from mitochondria that can prevent oxidation of mitochondrial DNA and subsequent DNA damage.

The MT200+Tr observed a tendency to higher values in most measured sperm characteristics than MT200 in the present study. Meanwhile, MT200+Tr, in contrast to MT200, indicated significantly greater motility, progressive motility, acrosome and membrane integrity, MMP, SCD, SOD, and TAC than MT20 and MT20+Tr. These positive results suggest the possibility that the MitoQ’s antioxidant action can have worked with the trehalose’s cryoprotective system in a synergy. However, elucidation of the mechanism of these synergistic effects needs further research.

Next, we also observed that if 2000 nM MitoQ (large dose) with or without trehalose is added to the extender, it cannot improve to the sperm function after cryopreservation. In line with our findings, several studies expressed that high concentrations of MitoQ (≥ 2 μM) in the semen extenders of human [20,21] and buffalo [6] increased ROS levels and a drop in mitochondrial potential. In this regard, Kumar et al. [20] have speculated that excessive MitoQ in mitochondria leads to increased concentration of conjugate decyl-TPP and subsequently impairs oxidative phosphorylation and mitochondrial function. Based on earlier and our data, it seems likely that the optimum dose of MitoQ in the freezing extender is essential to get the higher effectiveness associated with lower side effects in cryopreserved sperm quality of various animal species. Therefore, it is recommended that 200 nM MitoQ is considered an optimum antioxidant agent, and can be combined with trehalose in a Tris-based freezing extender of goat semen. However, further fertility studies are required to clarify sperm quality effects on the oocyte.

## 5. Conclusion

Supplementation of the extender with 200 nM MitoQ alone or combined with trehalose preserves the goat sperm quality characteristics and motion kinetics by preventing oxidative mitochondrial and DNA damage after cryopreserving. According to our findings, it appears that the MitoQ-induced reduction of oxidative sperm damage is mediated by reduced lipid peroxidation and improved antioxidant enzymatic defense; this can contribute to the cryostorage success of endangered Markhoz goat semen.

## Conflict of Interest

The authors report no conflict of interest.

## Acknowledgements

We are thankful to Dr. M. H. Sedri, Director of Kurdistan Agricultural and Natural Resources Research and Education Center, Sanandaj, Iran for providing the facility to conduct the research work. Sincere thanks to Dr. M. D. Joupari from Department of Animal and Marine Biotechnology, National Institute of Genetic Engineering and Biotechnology, Tehran, Iran for generously providing MitoQ. We must thank Mr. B. Rokhzad for assistance in experimental work.

## Notes

Funding: This research was no financial support.

### Competing Interest Statement

The authors have declared no competing interest.

## References

[1] E.M.E. Aboagla, Trehalose-Enhanced Fluidity of the Goat Sperm Membrane and Its Protection During Freezing, Biol. Reprod. 69 (2003) 1245–1250.

[2] S. Abril-Sánchez, A. Freitas-de-Melo, J. Giriboni, J. Santiago-Moreno, R. Ungerfeld, Sperm collection by electroejaculation in small ruminants: A review on welfare problems and alternative techniques, Anim. Reprod. Sci. 205 (2019) 1–9.

[3] H. Aebi, [13] Catalase in vitro, in: Oxyg. Radicals Biol. Syst., Academic Press, 1984: pp. 121–126.

[4] Z. Ahmad, M. Anzar, M. Shahab, N. Ahmad, S.M.H. Andrabi, Sephadex and sephadex ion-exchange filtration improves the quality and freezability of low-grade buffalo semen ejaculates, Theriogenology. 59 (2003) 1189–1202.

[5] F. Amidi, A. Pazhohan, M. Shabani Nashtaei, M. Khodarahmian, S. Nekoonam, The role of antioxidants in sperm freezing: a review, Cell Tissue Bank. 17 (2016) 745–756.

[6] V. Arjun, P. Kumar, R. Dutt, A. Kumar, R. Bala, N. Verma, A. Jerome, M. Virmani, C.S. Patil, S. Bhardwaj, D. Kumar, P.S. Yadav, Effect of mitochondria-targeted antioxidant on the regulation of the mitochondrial function of sperm during cryopreservation, Andrologia. (2022).

[7] N. Asadzadeh, Z. Abdollahi, S. Esmaeilkhanian, R. Masoudi, Fertility and flow cytometry evaluations of ram frozen semen in plant-based extender supplemented with Mito-TEMPO, Anim. Reprod. Sci. 233 (2021) 106836.

[8] M.N. Bucak, P.B. Tuncer, S. Sariözkan, P.A. Ulutaş, Comparison of the effects of glutamine and an amino acid solution on post-thawed ram sperm parameters, lipid peroxidation and anti-oxidant activities, Small Rumin. Res. 81 (2009) 13–17.

[9] D.R. Câmara, I. Ibanescu, M. Siuda, H. Bollwein, Mitoquinone does not improve sperm cryo-resistance in bulls, Reprod. Domest. Anim. 57 (2022) 10–18.

[10] K. Chandran, D. Aggarwal, R.Q. Migrino, J. Joseph, D. McAllister, E.A. Konorev, W.E. Antholine, J. Zielonka, S. Srinivasan, N.G. Avadhani, B. Kalyanaraman, Doxorubicin inactivates myocardial cytochrome c oxidase in rats: Cardioprotection by Mito-Q, Biophys. J. 96 (2009) 1388–1398.

[11] Z. Elyasi Gorji, P. Farzaneh, A. Nasimian, M. Ganjibakhsh, M. Izadpanah, M. Farghadan, F. Vakhshiteh, H. Rahmati, S.A. Shahzadeh Fazeli, H. Khaledi, A. Daneshvar Amoli, Cryopreservation of Iranian Markhoz goat fibroblast cells as an endangered national genetic resource., Mol. Biol. Rep. 48 (2021) 6241–6248.

[12] S. Escada-Rebelo, M.I. Cristo, J. Ramalho-Santos, S. Amaral, Mitochondria-Targeted Compounds to Assess and Improve Human Sperm Function, Antioxid. Redox Signal. 00 (2022).

[13] L. Fang, C. Bai, Y. Chen, J. Dai, Y. Xiang, X. Ji, C. Huang, Q. Dong, Inhibition of ROS production through mitochondria-targeted antioxidant and mitochondrial uncoupling increases post-thaw sperm viability in yellow catfish, Cryobiology. 69 (2014) 386–393.

[14] A. Farshad, Y. Hosseini, The cryoprotective effects of amino acids supplementation on cooled and post-thaw Markhoz bucks semen quality, Small Rumin. Res. 114 (2013) 258–263.

[15] J.L. Fernández, L. Muriel, M.T. Rivero, V. Goyanes, R. Vazquez, J.G. Alvarez, The sperm chromatin dispersion test: A simple method for the determination of sperm DNA fragmentation, J. Androl. 24 (2003) 59–66.

[16] J.F. Fonseca Torres, C.A.A., Maffili, V.V., Borges, A.M., Santos, A.D.F., Rodrigues, M.T., Oliveira, R.F.M., The hypoosmoctic swelling test in fresh goat spermatozoa, Anim. Reprod. 2 (2005) 139–144.

[17] K.A. Fortner, L.P. Blanco, I. Buskiewicz, N. Huang, P.C. Gibson, D.L. Cook, H.L. Pedersen, P.S.T. Yuen, M.P. Murphy, A. Perl, M.J. Kaplan, R.C. Budd, Targeting mitochondrial oxidative stress with MitoQ reduces NET formation and kidney disease in lupus-prone MRL-lpr mice, Lupus Sci. Med. 7 (2020).

[18] A. Garg, A. Kumaresan, M.R. Ansari, Effects of hydrogen peroxide (H 2O 2) on fresh and cryopreserved buffalo sperm functions during incubation at 37°C in vitro, Reprod. Domest. Anim. 44 (2009) 907–912.

[19] A.A. Ibrahim, H.M. Karam, E.A. Shaaban, M.M. Safar, M.F. El-Yamany, MitoQ ameliorates testicular damage induced by gamma irradiation in rats: Modulation of mitochondrial apoptosis and steroidogenesis, Life Sci. 232 (2019) 116655.

[20] P. Kumar, M. Wang, E. Isachenko, G. Rahimi, P. Mallmann, W. Wang, M. von Brandenstein, V. Isachenko, Unraveling Subcellular and Ultrastructural Changes During Vitrification of Human Spermatozoa: Effect of a Mitochondria-Targeted Antioxidant and a Permeable Cryoprotectant, Front. Cell Dev. Biol. 9 (2021) 1–20.

[21] L. Liu, M. Wang, T. Yu, Z. Cheng, M. Li, Q. Guo, [Mitochondria-targeted antioxidant Mitoquinone protects post-thaw human sperm against oxidative stress injury], Zhonghua Nan Ke Xue. 22 (2016) 205–211.

[22] C. Lv, G. Wu, Q. Hong, G. Quan, Spermatozoa Cryopreservation: State of Art and Future in Small Ruminants, Biopreserv. Biobank. 17 (2019) 171–182.

[23] G. Martin, N. Cagnon, O. Sabido, B. Sion, G. Grizard, P. Durand, R. Levy, Kinetics of occurrence of some features of apoptosis during the cryopreservation process of bovine spermatozoa, Hum. Reprod. 22 (2007) 380–388.

[24] T. Mitchell, D. Rotaru, H. Saba, R.A. Smith, M.P. Murphy, L.A. Mac-Millan-Crow, Correction to “The mitochondria-targeted antioxidant mitoquinone protects against cold storage injury of renal tubular cells and rat kidneys” (J Pharmacol Exp Ther (2011) 336, (682-692)), J. Pharmacol. Exp. Ther. 342 (2012) 596.

[25] N. Mohammadi, H. Daghigh Kia, G. Moghaddam, A. Javanmard, M. Murphy, Inhibition of ROS production by adding targeted antioxidants MitoQ and their effect on the performance of freeze-thawed rooster sperm, Anim. Sci. J. 32 (2019) 83–92.

[26] M.P. Murphy, R.A.J. Smith, Targeting antioxidants to mitochondria by conjugation to lipophilic cations, Annu. Rev. Pharmacol. Toxicol. 47 (2007) 629–656.

[27] A. Najafi, M. Zhandi, A. Towhidi, M. Sharafi, A. Akbari Sharif, M. Khodaei Motlagh, F. Martinez-Pastor, Trehalose and glycerol have a dose-dependent synergistic effect on the post-thawing quality of ram semen cryopreserved in a soybean lecithin-based extender, Cryobiology. 66 (2013) 275–282.

[28] N. Parajuli, L.H. Campbell, A. Marine, K.G.M. Brockbank, L.A. Macmillan-Crow, MitoQ blunts mitochondrial and renal damage during cold preservation of porcine kidneys., PLoS One. 7 (2012) 1–8.

[29] P. Peris-Frau, A.J. Soler, M. Iniesta-Cuerda, A. Martín-Maestro, I. Sánchez-Ajofrín, D.A. Medina-Chávez, M.R. Fernández-Santos, O. García-álvarez, A. Maroto-Morales, V. Montoro, J.J. Garde, Sperm cryodamage in ruminants: Understanding the molecular changes induced by the cryopreservation process to optimize sperm quality, Int. J. Mol. Sci. 21 (2020).

[30] A. Rezaei, A. Vaziry, A. Farshad, Letrozole, an aromatase inhibitor, improves seminal parameters and hormonal profile in aged endangered Markhoz bucks., Anim. Biosci. 00 (2021) 1–9.

[31] S. Schaäfer, A. Holzmann, The use of transmigration and SpermacTMstain to evaluate epididymal cat spermatozoa, Anim. Reprod. Sci. 59 (2000) 201–211.

[32] C. Schmelzer, F. Döring, Micronutrient special issue: Coenzyme Q10 requirements for DNA damage prevention, Mutat. Res. Mol. Mech. Mutagen. 733 (2012) 61–68.

[33] R.A.J. Smith, R.C. Hartley, M.P. Murphy, Mitochondria-targeted small molecule therapeutics and probes, Antioxidants Redox Signal. 15 (2011) 3021–3038.

[34] R.A.J. Smith, M.P. Murphy, Animal and human studies with the mitochondria-targeted antioxidant MitoQ, Ann. N. Y. Acad. Sci. 1201 (2010) 96–103.

[35] D.F. Stowe, A.K.S. Camara, Mitochondrial reactive oxygen species production in excitable cells: Modulators of mitochondrial and cell function, Antioxidants Redox Signal. 11 (2009) 1373–1414.

[36] Y. Sui, Q. Fan, B. Wang, J. Wang, Q. Chang, Ice-free cryopreservation of heart valve tissue: The effect of adding MitoQ to a VS83 formulation and its influence on mitochondrial dynamics, Cryobiology. 81 (2018) 153–159.

[37] S. Tiwari, R.K. Dewry, R. Srivastava, S. Nath, T.K. Mohanty, Targeted antioxidant delivery modulates mitochondrial functions, ameliorates oxidative stress and preserve sperm quality during cryopreservation, Theriogenology. 179 (2022) 22–31.

[38] S. Tiwari, T.K. Mohanty, M. Bhakat, N. Kumar, R.K. Baithalu, S. Nath, H.P. Yadav, R.K. Dewry, Comparative evidence support better antioxidant efficacy of mitochondrial-targeted (Mitoquinone) than cytosolic (Resveratrol) antioxidant in improving in-vitro sperm functions of cryopreserved buffalo (Bubalus bubalis) semen, Cryobiology. 101 (2021) 125–134.

[39] W.Y. Wani, S. Gudup, A. Sunkaria, A. Bal, P.P. Singh, R.J.L. Kandimalla, D.R. Sharma, K.D. Gill, Protective efficacy of mitochondrial targeted antioxidant MitoQ against dichlorvos induced oxidative stress and cell death in rat brain, Neuropharmacology. 61 (2011) 1193–1201.

[40] C. Wu, J. Dai, S. Zhang, L. Sun, Y. Liu, D. Zhang, Effect of Thawing Rates and Antioxidants on Semen Cryopreservation in Hu Sheep, 19 (2021) 204–209.

[41] C. Wu, J. Dai, S. Zhang, L. Sun, Y. Liu, D. Zhang, Effect of Thawing Rates and Antioxidants on Semen Cryopreservation in Hu Sheep, Biopreserv. Biobank. 19 (2021) 204–209.

[42] Y. Wu, C. Hao, X. Liu, G. Han, J. Yin, Z. Zou, J. Zhou, C. Xu, MitoQ protects against liver injury induced by severe burn plus delayed resuscitation by suppressing the mtDNA-NLRP3 axis, Int. Immunopharmacol. 80 (2020) 106189.

[43] I. Yánez-Ortiz, J. Catalán, J.E. Rodríguez-Gil, J. Miró, M. Yeste, Advances in sperm cryopreservation in farm animals: Cattle, horse, pig and sheep, Anim. Reprod. Sci. (2021) 1–18.

[44] J. Zhang, X. Bao, M. Zhang, Z. Zhu, L. Zhou, Q. Chen, Q. Zhang, B. Ma, MitoQ ameliorates testis injury from oxidative attack by repairing mitochondria and promoting the Keap1-Nrf2 pathway, Toxicol. Appl. Pharmacol. 370 (2019) 78–92.

